# Benchtop micro-mashing: high-throughput, robust, experimental beer brewing

**DOI:** 10.1101/2020.06.24.158345

**Authors:** Edward D. Kerr, Christopher H. Caboche, Peter Josh, Benjamin L. Schulz

## Abstract

Brewing science is undergoing a renaissance with the use of modern analytical chemistry and microbiology techniques. However, these modern analytical tools and techniques are not necessarily aligned with the scale and scope of brewing science. In particular, brewing processes can be time consuming, ingredient intensive, and require specialised technical equipment. These drawbacks compound with the need for appropriate numbers of replicates for adequately powered experimental design. Here, we describe a micro-scale mash method that can be performed using a common laboratory benchtop shaker/incubator, allowing for high throughput mashing and easy sample replication for statistical analysis. Proteomic profiles at both the protein and peptide levels were consistent between the 1 mL micro-mash and a 23 L Braumeister mash, and both mash scales produced wort with equivalent fermentable sugar and free amino acid profiles. The experimental flexibility offered by our micro-mash method allowed us to investigate the effects of altered mash parameters on the beer brewing proteome.

## Introduction

Beer brewing is one of the oldest known biotechnologies and has heavily influenced society and science. The ancient science of beer brewing is rapidly developing, with modern analytical techniques allowing deeper understanding of fundamental aspects of its integrated bioprocesses^1–10^. The brewing process begins with harvested barley seeds being malted through a process which involves controlled, partial germination followed by kilning^11,12^. The malt is then milled to open the husks and combined with warm water in a process known as mashing. Mashing extracts the soluble sugars and proteins from the malt which are digested by enzymes into smaller sugars and free amino nitrogen (FAN)^11^. Wort, the soluble fraction of the mash, is then boiled with hops to introduce bitterness and aromatic flavours, remove volatile off-flavours, stop enzymatic activity, and sterilize^11,13^. Fermentation is performed by pitching yeast into cooled wort, where the yeast consume the sugars and FAN and produce flavour compounds, carbon dioxide, and ethanol. Finally, beer is filtered, aged, carbonated, and packaged.

Complex and dynamic protein and carbohydrate chemistry occur during mashing and the boil^1^. These processes are very well understood from an industrial standpoint, but the underlying biochemical details are only starting to be appreciated^11,12^. Mashing extracts the soluble component from malted barley seeds, primarily starch and proteins, including enzymes^11^. Extracted starch and proteins are then broken down by enzymes into small fermentable sugars and FAN, respectively^11^. Proteomics has been used to identify and monitor changes in the abundance of these enzymes and other proteins during the mash and boil and into the final beer product^1,4–10^. These studies found that while some proteins were heat-stable and could survive being boiled, other proteins were only active and stable at low temperatures, while many proteins were covalently modified by a variety of processes during brewing^1,4–10,14^.

Brewing processes at the scale of a large industrial brewery or even a smaller craft brewery are time consuming, ingredient intensive, and require specialised technical equipment not typically found in research laboratories. This large scale makes it difficult and expensive to perform appropriately designed experiments with sufficient statistical power to advise the brewing process.

Our previous investigation of the beer brewing proteome identified and measured the abundance of hundreds of proteins during the mash, and found dynamic changes in protein abundance due to solubilisation, proteolysis, and degradation^1^. Here, we re-interrogate these datasets from a 23 L mash and boil and a 1 mL benchtop micro-mash^1^. We investigate the two datasets in more detail to test the reliability and comparability of a 1 mL benchtop micro-mash to large scale brewing. We present a micro-scale mash protocol that can be performed using a benchtop shaker incubator, allowing for high-throughput mashing and easy sample replication for statistical analysis (Fig. 1).

**Figure 1.**
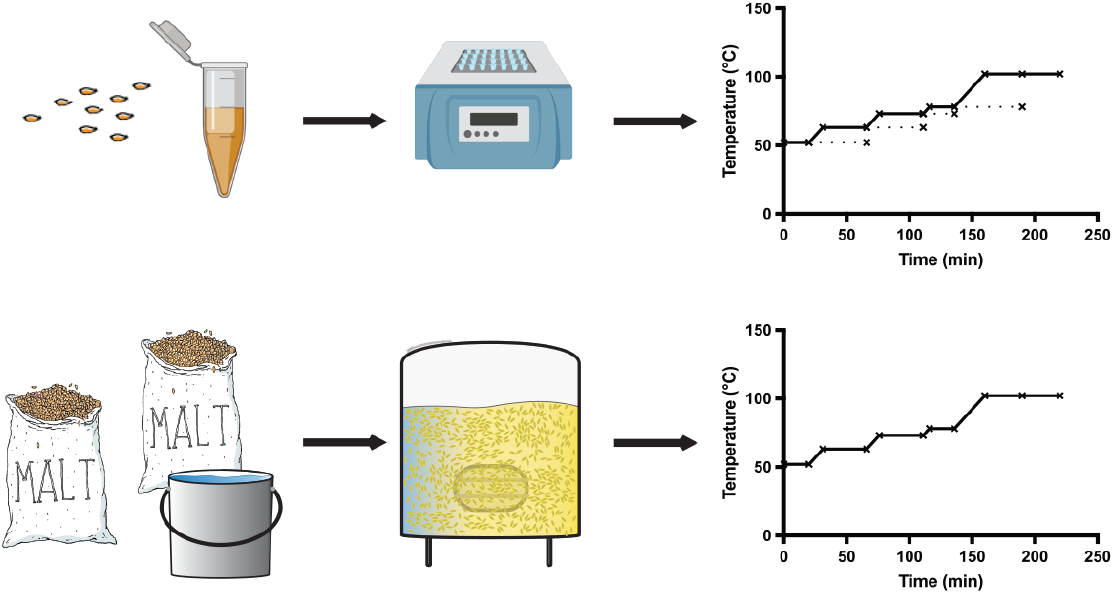
Process and throughput of micro-scale mash compared with a standard Braumeister mash.

## Results

### Protein abundance profiles are similar in 23 L Braumeister and 1 mL benchtop mashes

We interrogated the abundance profile of each individual protein from a 23 L Braumeister mash and a 1 mL benchtop mash (Fig. 2A). These protein abundance profiles showed that at both mash scales proteins increased in abundance as temperature increased and proteins were solubilised. As temperature continued to increase, the temperature-sensitive proteins reduced in abundance due to denaturation, aggregation, and precipitation (Fig. 2A). Some subtle differences are apparent between the 23 L and 1 mL mash scales, with protein extraction being slightly faster in the 1 mL micro-scale mash than in the 23 L Braumeister mash (Fig. 2A). However, the profile of a given protein was generally very similar at both 23 L and 1 mL mash scales, while the profiles for different proteins were distinct.

**Figure 2.**
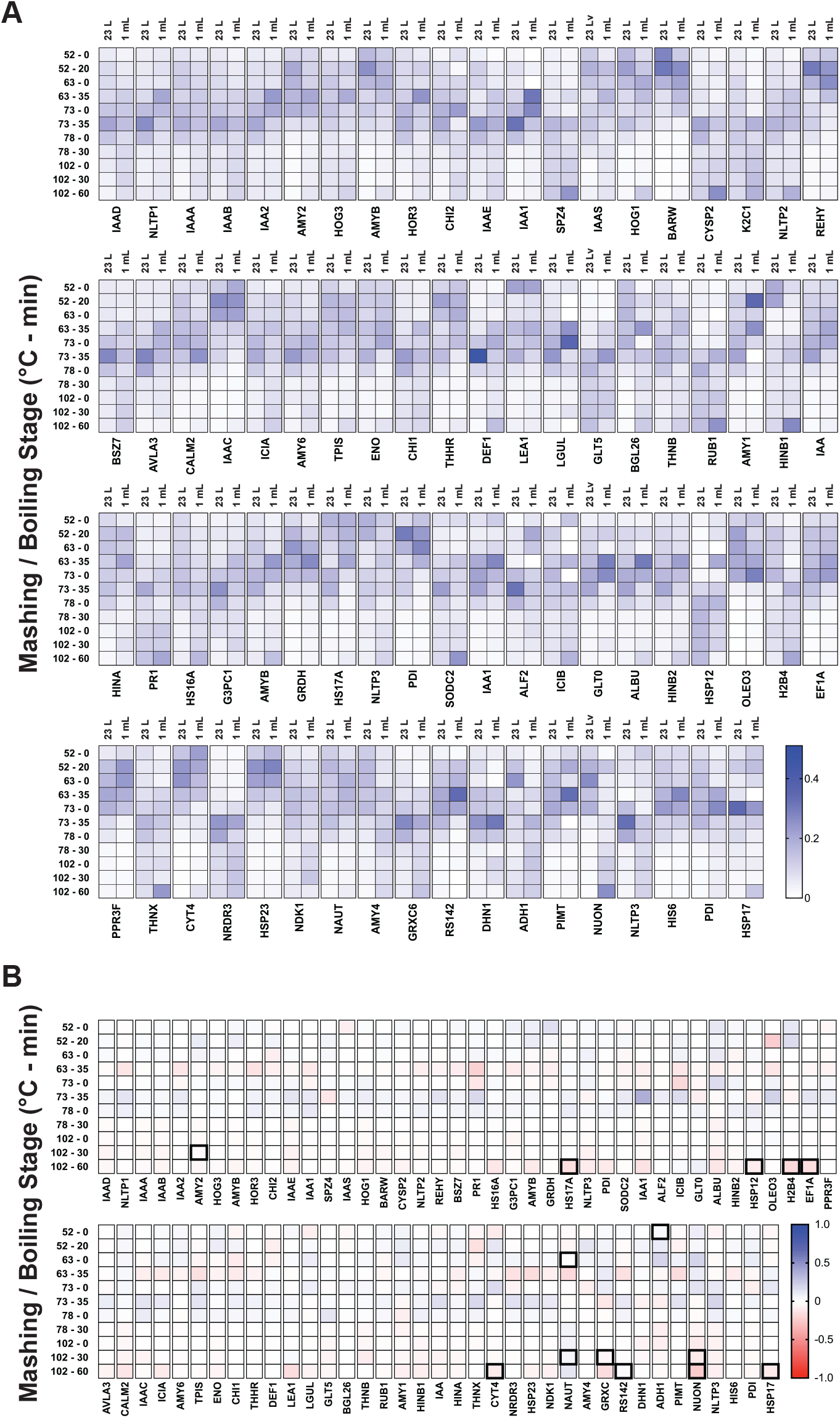
Overall protein abundance profiles are similar between a 23 L Braumeister mash and a 1 mL micro-scale mash. **(A)** Normalised abundance at each time point in 23 L Braumeister and 1 mL micro-scale mashes, for all proteins identified. Relative protein abundance was calculated as protein intensity at each point relative to trypsin intensity at the same point. Normalised abundance was then calculated as the relative protein abundance at each point relative to the combined relative protein abundance across all points. **(B)** Difference of the normalised protein abundance means at each time point between 23 L Braumeister and 1 mL micro-scale mashes. Positive values (blue), higher in 23 L Braumeister; negative values (red), higher in 1 mL micro-scale. Bold highlights, statistically significant (p < 0.05).

We next directly compared the performance of the two mash scales by calculating the difference in normalized abundance, subtracting the normalized abundance of each protein at each stage between the two mashes (Fig. 2B and Supplementary Table S1). This difference map was consistent with the overall protein abundance profiles (Fig. 2A) and demonstrated that proteins were more efficiently extracted in the 1 mL micro-scale mash, with increased levels at the end (35 min) of the 63 °C rest, although these differences were not statistically significant (Fig. 2B). This difference map also demonstrated that the loss of protein at higher temperatures was less efficient in the 1 mL micro-scale mash than in the 23 L Braumeister mash (Fig. 2B). This difference was particularly apparent during the boil. Regardless of these minor differences in the rates of extraction and denaturation, these side-by-side and difference comparisons of the two mash scales showed their protein-specific abundance profiles were very similar overall, with very few statistically significant differences (Fig. 2).

### Proteolytically cleaved forms of peptides show similar profiles in a 23 L Braumeister mash and a 1 mL benchtop mash

Proteolysis of barley seed storage proteins releases FAN for yeast to consume during fermentation and has also been shown to be a key driver of protein stability during mashing^1,11^. Barley seeds contain proteases which can proteolytically clip barley seed storage proteins. We previously found that in a 23 L Braumeister mash, the proteolytically clipped forms of proteins tend to be less stable at higher temperatures in the mash, which results in loss of these non-tryptic peptides and an increase in the relative abundance of the full-length tryptic peptides^1^. However, this effect is site-specific, as some proteolytically clipped forms show high stability. We therefore compared this key feature of the dynamic mashing proteome at 23 L Braumeister and 1 mL micro mash scales. This analysis showed that the abundance of site-specific proteolysis products was very similar between the mash scales, with only slight differences detected at lower temperatures early in the mash (Fig. 3A and B).

**Figure 3.**
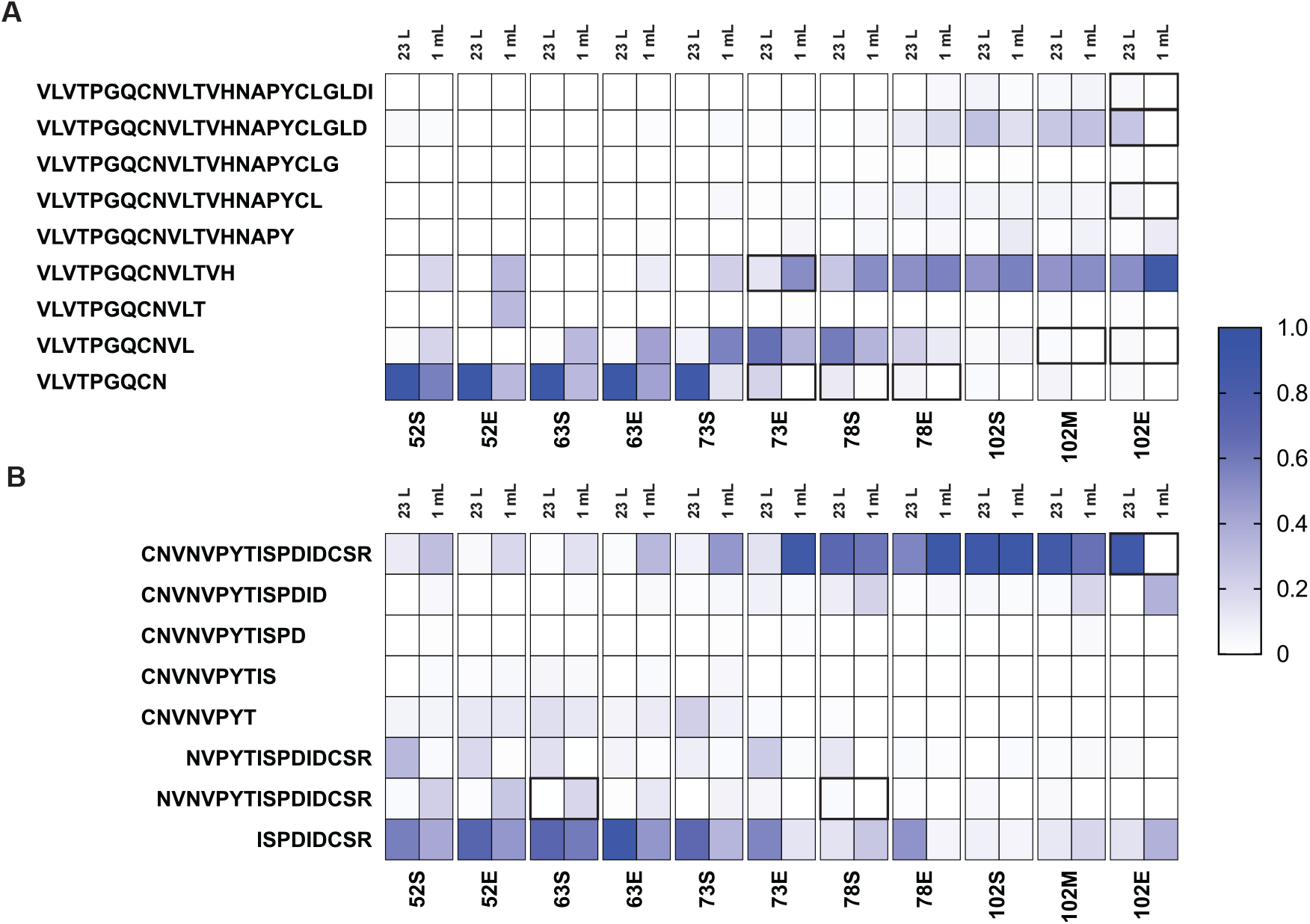
Proteolytically clipped peptide abundance profiles are similar between a 23 L Braumeister and a 1 mL micro-scale mash. **(A)** Normalised abundance of full and semi-tryptic peptides of V_122_LVTPGQCNVLTVHNAPYCLGLDI_145_ from IAAA in both the Braumeister mash (left columns) and micro-scale mash (right columns). **(B)** Normalised abundance of full and semi-tryptic peptides of K-C_99_NVNVPYTISPDIDCSR_115_-I from NLTP1 in both the Braumeister mash (left columns) and benchtop micro-scale mash (right columns). Values show mean, n=3. Bold highlights, statistically significant (p < 0.05).

For proteolytic forms of the peptide family V_122_LVTPGQCNVLTVHNAPYCLGLDI_145_ from α-amylase/trypsin inhibitor CMa (IAAA) we found almost identical peptide abundance profiles between the two mash scales, with few statistically significant differences (Fig. 3 and Supplementary Table S2). The semi-tryptic forms V_122_LVTPGQCNVLTVH_135_ and V_122_LVTPGQCNVL_132_ were slightly increased in abundance at lower temperatures in the 1 mL mash compared to the 23 L Braumeister mash, but these differences converged at higher temperatures later in the mash (Fig. 3A). We also found a high level of similarity in peptide abundance profiles between the two mash scales for proteolytic forms from the peptide family K-C_99_NVNVPYTISPDIDCSR_115_-I from non-specific lipid-transfer protein 1 (NLTP1) (Fig. 3B). Full tryptic peptide K-C_99_NVNVPYTISPDIDCSR_115_-I and the semi-tryptic form N_100_VNVPYTISPDIDCSR_115_-I were slightly higher in abundance early in the 1 mL micro-mash, but again these differences decreased as the mash progressed (Fig. 3B). At both mash scales, both of these peptide groups showed equivalent overall profiles when comparing the dynamics of proteolysis and the resulting protein stability across the mash and boil (Fig. 3).

### Statistical power of benchtop micro-scale mashing

A major difficulty in performing rigorous experiments using brewing equipment at an industrial scale is the cost in time and reagents of performing sufficient numbers of replicates to allow statistical comparisons. We have shown that the performance of a micro-mash at a 1 mL scale was equivalent to that of a large scale 23 L Braumeister mash in terms of protein (Fig. 2) and peptide (Fig. 3) abundance profiles during the mash and boil. The small scale of the micro-mash allows it to be easily used to perform experimental manipulation of mash parameters, such as the time and temperature of each rest, while increasing the number of samples to allow for the replicates and controls needed for statistical comparisons. The ability to control and alter the time and temperature of each rest (stage) in the mash allows for distinction between time- and temperature-dependent changes in protein and peptide abundance (Fig. 4). This can be coupled with statistical analysis to test whether these changes are indeed time- or temperature-dependent (Fig. 4).

**Figure 4.**
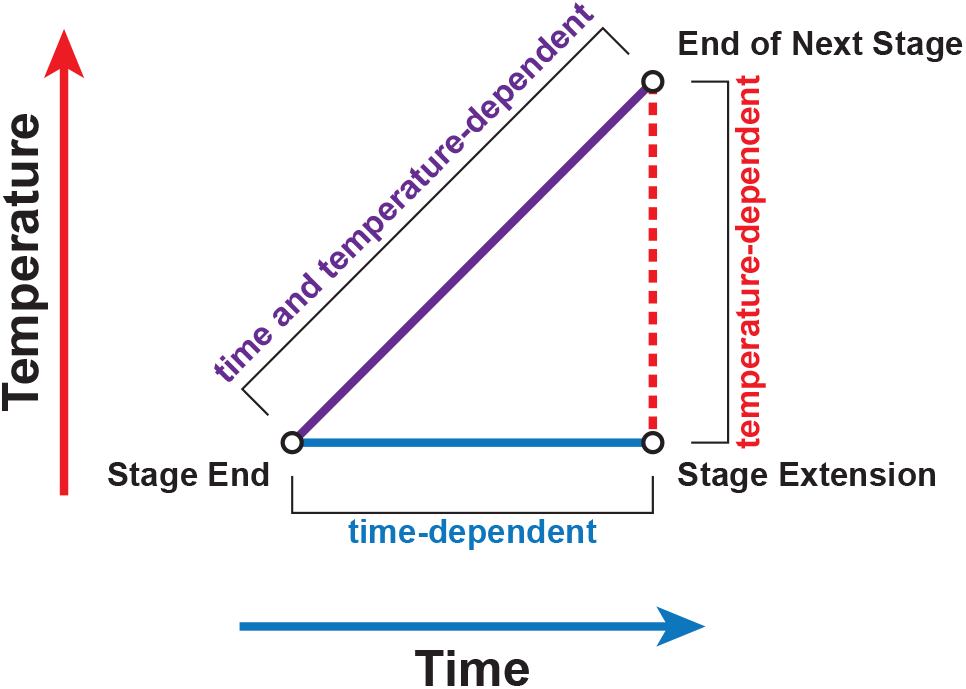
Mash stage extension allows differentiation between time- and temperature-dependent changes in protein abundance. Statistical comparison in protein or peptide abundance between the end of a stage, extension of the stage, and end of the next stage allows the determination of the cause of changes in abundance. A significant difference between ‘stage end’ and ‘stage extension’ corresponds to a time-dependent change in protein abundance. A significant difference between ‘end of next stage’ and ‘stage extension’ corresponds to a temperature-dependent change in protein abundance. A significant difference between only ‘stage end’ and ‘end of next stage’ corresponds to a time- and temperature-dependent change in protein abundance.

To determine if protein changes during the mash were time- or temperature-dependent, we previously performed a micro-scale mash that included extended incubation periods at each temperature stage^1^. This analysis showed that the cause of changes in abundance (time- or temperature-dependent) could be established for both NLTP1 and IAAA using these temperature stage extensions (Fig. 5).

**Figure 5.**
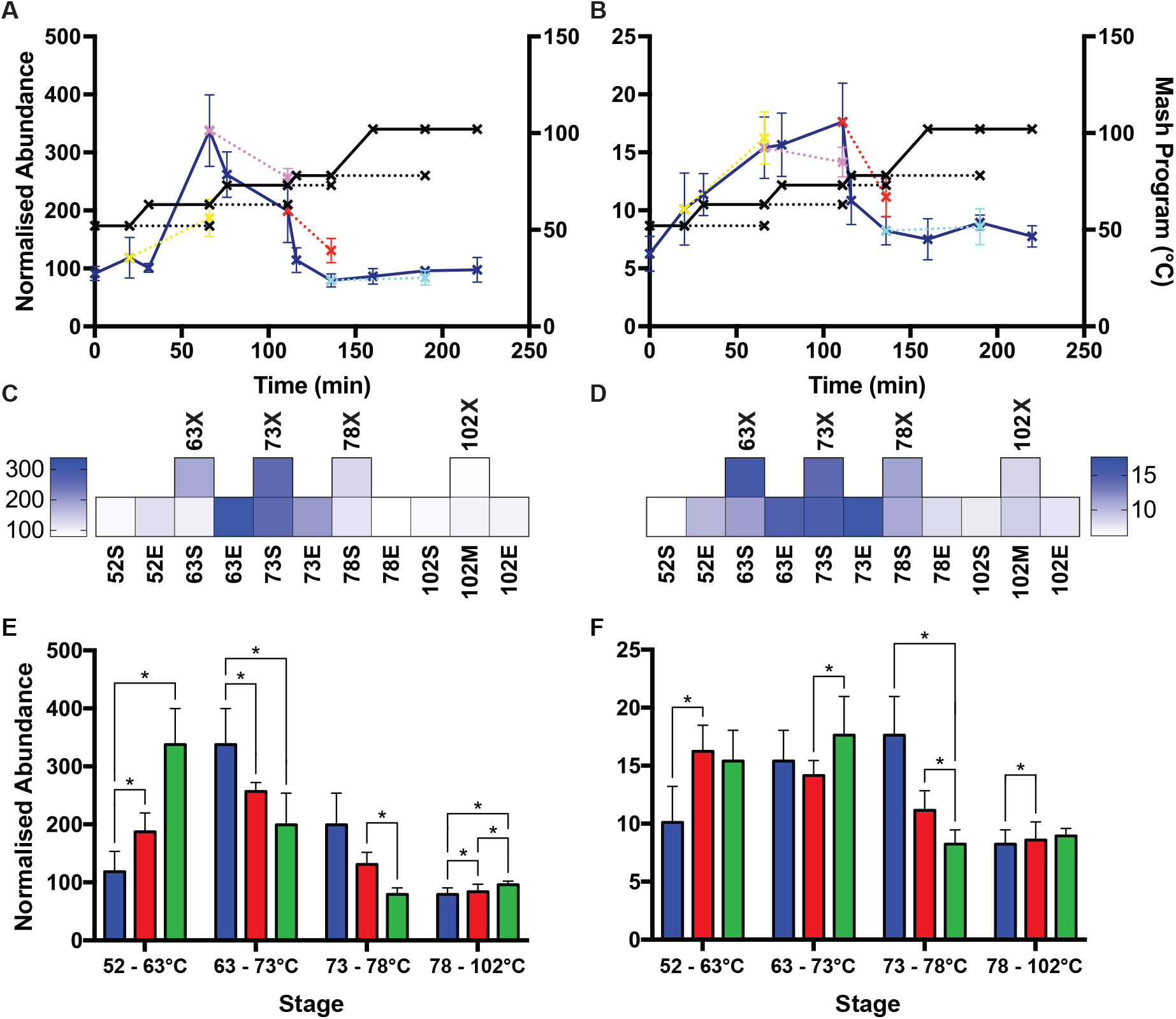
The effect of time and temperature on protein abundance during the mash and boil in a 1 mL micro-scale mash. Normalised abundance of (**A**) NLTP1 and **(B)** IAAA. Standard mash program, solid line, dark blue; rest stage temperature extensions, dotted lines. Heat map of normalised abundance of (**C**) NLTP1 and **(D)** IAAA. Standard mash program, bottom row; rest stage temperature extensions, top row. Comparison of normalised abundance between sequential stages for (**E**) NLTP1 and (**F**) IAAA). First stage, blue; time extension of first stage, red; next temperature stage, green. All values show mean, n=3. Error bars, SEM. *, statistically significant (p < 0.05).

### Discovering if changes to peptide abundance are time- or temperature-dependent using mash stage extensions

Proteolysis is a key post translational modification in brewing, driving FAN production and controlling protein stability^11^. We previously found that many physiological proteolysis events could be measured during the mash, and that the relative abundance of the corresponding differently proteolytically clipped protein forms changed over the course of the mash^1^. The observed changes were consistent with limited proteolysis occurring in malting or early in the mash, and the subsequent proteolytic clipping affecting protein stability by altering the temperature at which the protein unfolded, aggregated, and was lost from solution. Most proteolytic events lowered protein stability, while a few had no effect or even increased stability. To investigate the mechanisms underlying these effects in more detail, we studied the effect of the temperature stage extensions on the abundance of peptides resulting from physiological proteolysis. For two highly abundant peptide families, V_122_LVTPGQCNVLTVHNAPYCLGLDI_145_ from IAAA and K-C_99_NVNVPYTISPDIDCSR_115_-I from NLTP1, the relative abundance of each peptide form was statistically compared between stages of the mash (Fig. 6C and F, and Supplementary Tables S3 and S4).

**Figure 6.**
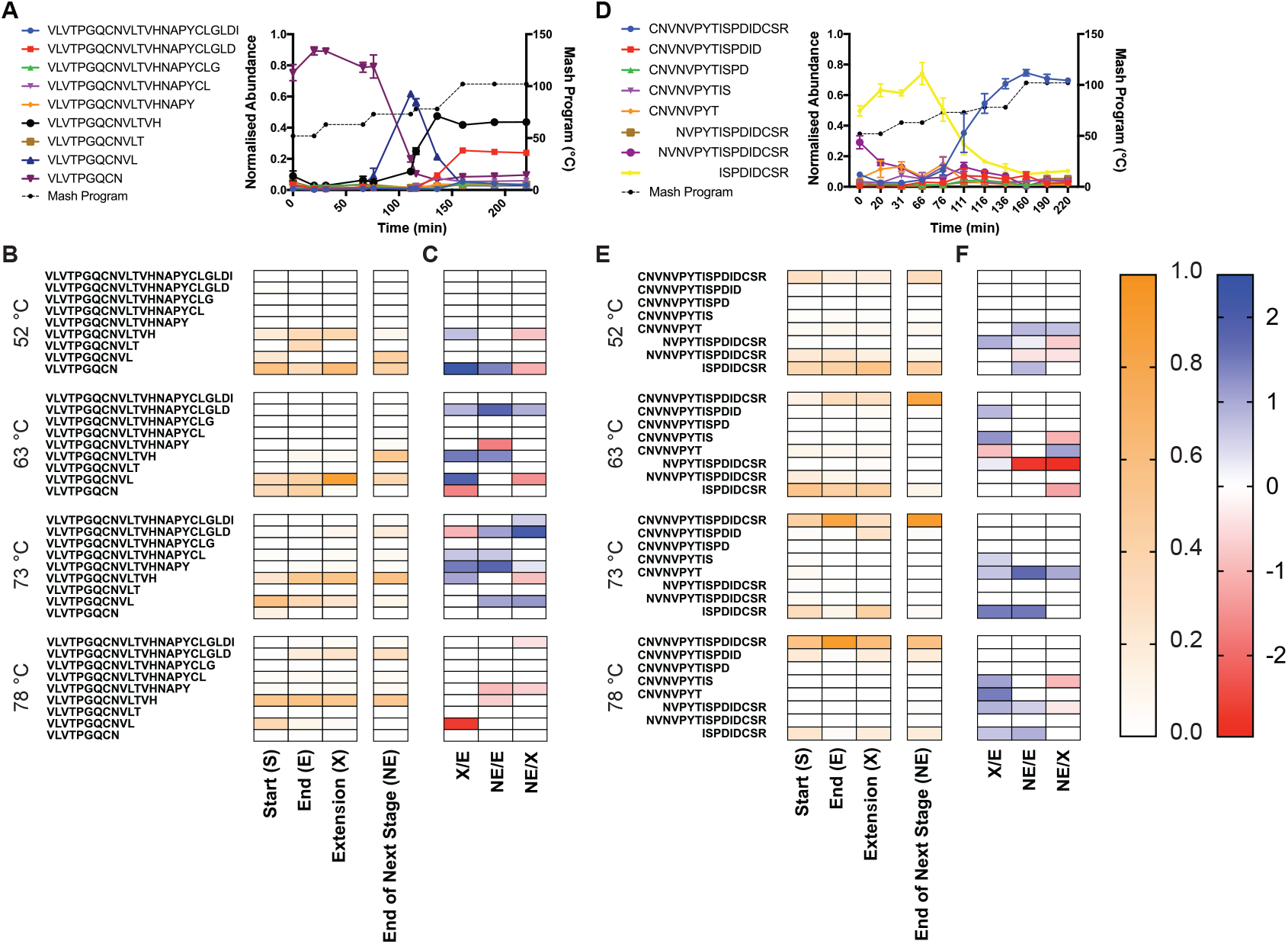
Time- and temperature-dependent abundance of full- and semi-tryptic peptides during the mash and boil. **(A)** Normalised abundance (a.u; arbitrary units) in a 23 L mash of full- and semi-tryptic peptides corresponding to V_122_LVTPGQCNVLTVHNAPYCLGLDI_145_ from IAAA**. (B)** Heatmap of normalised abundance in a 1 mL mash of IAAA full- and semi-tryptic peptides corresponding to V_122_LVTPGQCNVLTVHNAPYCLGLDI_145_ at the start, end, extension, and end of the next step across the mash. **(C)** Log_2_ (fold change) of peptides that were significantly differentially abundant (p < 10^−5^) between the designated stages of the 1 mL mash: end of stage (E), extension of stage (X), end of the next stage (NE). **(D)** Normalised abundance (a.u; arbitrary units) in a 23 L mash of full- and semi-tryptic peptides corresponding to C_99_NVNVPYTISPDIDCSR_115_-I from NTLP1**. (E)** Heatmap of normalised abundance in a 1 mL Mash of NLTP1 full- and semi-tryptic peptides corresponding to K-C_99_NVNVPYTISPDIDCSR_115_-I at the start, end, extension, and end of the next step across the mash. **(F)** Log_2_ (fold change) of peptides that were significantly differentially abundant (p < 10^−5^) between the designated stages of the 1 mL mash: end of stage (E), extension of stage (X), end of the next stage (NE). (A and D) Values show mean, n=3. Error bars show SEM. Mash temperature profile is shown on the right Y-axis. Mash program is shown in black dotted lines; points indicate when samples were taken. (B and E) Values show mean, n=3.

For proteolytic forms of the peptide family V_122_LVTPGQCNVLTVHNAPYCLGLDI_145_, we observed that some smaller semi-tryptic forms were more abundant than the full tryptic peptide at lower temperatures (52 °C and 63 °C), and that at higher temperatures two semi-tryptic forms increased in relative abundance (Fig. 6A and B), as previously described^1^. We found that for the semi-tryptic peptide V_122_LVTPGQCN, the decrease in abundance after the 63 °C rest was time-dependent (Fig. 6A, B, and C). The slightly larger form V_122_LVTPGQCNL increased in relative abundance at 73 °C, which was temperature-dependent (Fig. 6A, B, and C). At the 78 °C rest, a sharp decrease in abundance of V_122_LVTPGQCNL occurred, which was time-dependent (Fig. 6A, B, and C). Finally, during the 78 °C rest the relative abundance of bothV_122_LVTPGQCNVLTVHNAPYCLGLD and V_122_LVTPGQCNVLTVH increased (Fig. 6A, B, and C). The increase of V_122_LVTPGQCNVLTVHNAPYCLGLD was temperature-dependent, while the increase of V_122_LVTPGQCNVLTVH was time-dependent (Fig. 6A, B, and C).

For proteolytic forms of the peptide family C_99_NVNVPYTISPDIDCSR_115_-I, we observed that smaller semi-tryptic forms, predominantly ISPDIDCSR_115_-I. were more abundant than the full tryptic peptide at lower temperatures (52 °C and 63 °C), and that at higher temperatures the full tryptic peptide was the predominant form (Fig. 6D and E), as previously described^1^. We found that for the semi-tryptic peptide ISPDIDCSR_115_-I, the increase in abundance after the 52 °C rest was temperature-dependent (Fig. 6D, E, and F). Following this, ISPDIDCSR_115_-I increased in relative abundance at 63 °C, which was temperature-dependent (Fig. 6D, E, and F). The full tryptic form C_99_NVNVPYTISPDIDCSR_115_-I appeared to increase in abundance at the 73 °C rest, but no comparison was significant (Fig. 6D, E, and F).

### Wort metabolomic analysis

The micro-mash system closely mirrors the performance of a 23 L Braumeister mash when considering the dynamic proteome of the mash and boil. As well as the proteome, other key parameters relevant to mashing performance are the specific gravity, FAN or soluble amino acid levels, and sugar profiles of the final wort produced by the mashing process. We compared these parameters for the 1 L micro-scale and 23 L Braumeister mashes. Multiple Reaction Monitoring (MRM) LC-MS/MS analysis showed that the fermentable sugar profiles after the mash and boil using the two different mash methods were very similar, with no significant differences (Fig. 7A), while the wort from the 1 mL micro-scale mash had higher specific gravity (Fig. 7B). Finally, we used LC-MS/MS to measure 15 free amino acids, which showed that wort from the 1 mL micro-scale mash contained very similar concentrations of free amino acids as the 23 L Braumeister mash, with no significant differences (Fig 7C).

**Figure 7.**
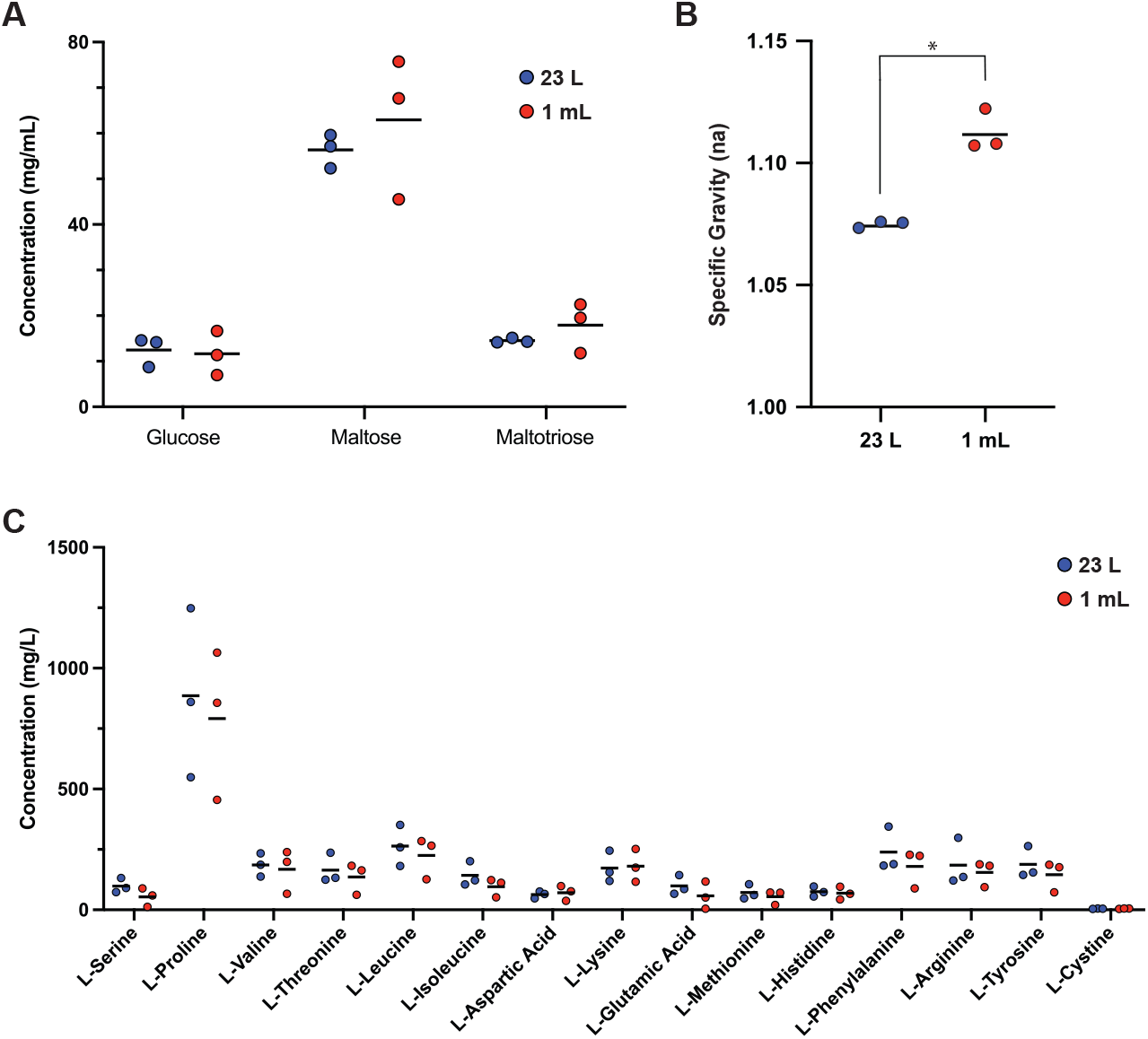
Wort parameters from a 23 L Braumeister mash and a 1 mL micro-scale mash. **(A)** Glucose, maltose, and maltotriose quantified in wort from a 23 L Braumeister mash (blue) and a 1 mL micro-mash (red). (**B**) Specific gravity. **(C)** Amino acids quantified in the two mashes. Bars show mean, n=3. *, statistically significant (p < 0.05).

## Discussion

We have demonstrated that a benchtop micro-mashing method can be used in lieu of standard industrial-scale brewing equipment to investigate the molecular dynamics of the brewing process. Specifically, a 1 mL micro-scale mash was not only equivalent to a full 23 L Braumeister mash but allowed increased throughput for statistical analyses. We showed that the protein abundance profiles of the 23 L Braumeister and 1 mL micro-scale mashes were highly similar (Fig. 2). The main difference between the mash scales was an increased protein abundance earlier in the 1 mL micro-scale mash compared to the 23 L Braumeister mash, consistent with rapid extraction and solubilization of proteins in the micro-mash. (Fig. 2). We found that the abundance of proteolysis products in the form of semi-tryptic peptides and their full-length tryptic counterparts were very similar between the two mashes. There were only small differences between mash scales, with proteolysis products being more abundant at the lower temperatures in the micro-mash (Fig. 3).

The temperature stage extensions coupled with statistical tests provided a model for statistically determining if changes to protein or peptide abundance during the mash were temperature- and/or time-dependent^1^. At the protein level, stage extensions provided a simple tool to determine the cause of changes in protein abundance (Fig. 5). When investigating peptide abundance and proteolysis, temperature controls provided insight into how proteolysis affected peptide abundance and protein stability (Fig. 6). Our results here showed that coupling mash stage extensions with replicate analyses and statistical tests allows for insight into the environmental and process causes of changes in the abundance and modification of proteins. Our data showed that protein and proteolysis biochemistry in the mash is highly complex, and tools such as this micro-mash method will allow future interrogation of this complex biochemistry.

We found that the 1 mL micro-scale and 23 L Braumeister mashes produced wort with very similar concentrations of fermentable sugars and free amino acids. Glucose, maltose, and maltotriose as well as all the measured amino acids had equivalent concentrations at both mash scales (Fig. 7). The final specific gravity was higher in the 1 mL mico-scale mash (Fig. 7), possibly because of more efficient solubilization of starch or maltooligosacharides at this scale. The micro-mash method we describe here provides a high-throughput and low-cost tool to study brewing biochemistry. The temperature extensions made possible by the micro-scale and high-throughput system provide a novel and highly practical tool to systematically unravel changes in the abundance of proteins and their post-translational modifications. This micro-mash system will also allow high-throughput analysis of barley varieties that are being accredited or analysed commercially.

## Methods

### Braumeister Mash and Benchtop Micro-Mash

Previously published^1^ mass spectrometry proteomics datasets PXD010013 and PXD013177 were retrieved from the ProteomeXchange Consortium via the PRIDE partner repository^15^. The mash methods that generated the datasets were previously described^1^. Briefly, a mash was performed using a Braumeister (Speidel) with 23 L of water and 5 kg of milled Vienna pale ale malt with a multi-step mash. Samples were taken at the start and end of each stage, and after 30 min of boil. Samples were taken from independent triplicate mashes using malt from the same batch. Micro-scale mashing was performed using the same multi-step mash profile as the Braumeister mash with additional time extensions at each stage of the mash. Each replicate sample was prepared individually in a 1.5 mL protein LoBind tube (Eppendorf) with 200 mg of milled pale ale malt resuspended in 1.0 mL of water. Samples were incubated in a Multi-Therm Heat-Shaker incubator (Benchmark Scientific) shaking at 1500 rpm with manual temperature control. Before the boil, each sample was clarified by centrifugation at 18,000 rcf for 1 min at room temperature, the supernatant transferred to a new protein LoBind tube, and incubated with an open lid on a static heat block preheated to 102°C. Samples were taken at the start, end, and extension of each stage, and after 30 min of boil. Samples were immediately clarified by centrifugation at 18,000 rcf for 1 min at room temperature, and the supernatants were transferred to new Protein LoBind tubes and stored at −20°C.

### Sample Preparation and Mass Spectrometry

Sample preparation for previously published^1^ mass spectrometry proteomics datasets PXD010013 and PXD013177 included methanol/acetone precipitation of proteins from 10 μL or 100 μL of each sample from Braumeister- or micro-scale mashes, respectively, followed by digestion with trypsin in the presence of 10 mM dithiothreitol. After desalting with C18 ZipTips, peptides were measured with Data Dependent Acquisition and Data Independent Acquisition LC-ESI-MS/MS as previously described^1,16^.

### Data Analysis

Proteins and peptides were identified with Protein Pilot 5.1 (SCIEX) as previously described^1,16^. The abundance of peptide fragments, peptides, and proteins was determined using PeakView 2.1 (SCIEX), with settings: shared peptides, allowed; peptide confidence threshold, 99%; false discovery rate, 1%; XIC extraction window, 6 min; XIC width, 75 ppm. PeakView 2.1 processing for abundances were performed independently for Braumeister and benchtop micro-mash both using the ion library created for previous Braumeister mash^1^. For protein-centric analyses, protein abundances were normalised to the abundance of trypsin self-digest peptides, as previously described^1^. For comparison of protein abundance between mashes, the trypsin-normalised abundance of each protein at each time point was normalised to the summed trypsin-normalised abundance of that protein across all time points. Differences at each point between mashes were determined by Student’s t-test with Bonferroni correction for multiple comparisons and a significance threshold of p = 0.05. For analysis of peptide modifications, the abundance of each form within a peptide family was normalised to the summed abundance for all detected forms within the peptide family. For determining time- and temperature-dependent changes, statistical analysis was done by reformatting the PeakView output using python script (https://github.com/bschulzlab/reformatMS)^17^, and significant differences in protein and peptide abundances between stages were determined using MSstats (2.4) in R (21), with a significance threshold of p = 0.05 as previously implemented^1^.

### Sugar and Amino Acid Quantification by MRM LC-MS/MS

Samples were filtered with a 30 kDa Amicon Ultra centrifuge filter unit (Millipore) and diluted 1:1000 with H_2_O. Sugars and amino acids were measured by MRM with UPLC-MS/MS using a Nexera ultraperformance LC system (Shimadzu) coupled to a QTRAP 5500 mass spectrometer (SCIEX). Chromatography was performed with a Kinetex 1.7 μm C18 100 mm x 2.1 mm column (Phenomenex) in a 50 °C column oven. The MS was operated in negative MRM mode for sugars with source parameters: ISV, −4500 V; temperature, 400 °C; curtain gas, 20; GS1, 45; and GS2, 50. Amino acid analysis was performed in positive MRM mode with source parameters: ISV, 5500 V; temperature, 400 °C; curtain gas, 20; GS1, 50; and GS2, 60. For sugar analysis, an isocratic elution program at a flow rate of 0.4 mL/min was used, with buffer A (1 % acetonitrile and 0.1% formic acid) and buffer B (90% acetonitrile and 0.1% formic acid) (0 min, 0% B; 5 min, 0% B; 5:10 min, 100% B; 10 min, 100% B) for a total run time of 10 min. For amino acid analysis, a gradient elution program at a flow rate of 0.4 mL/min was used, with buffer A (1 % acetonitrile and 0.1% formic acid) and buffer B (90% acetonitrile and 0.1% formic acid) (0 min, 1% B; 1 min, 1% B; 5 min, 20% B; 5:50 min, 100% B; 6:50 min, 100% B; 7 min, 1% B; 10 min, 1% B) for a total run time of 10 min. Sugars (glucose, maltose, and maltotriose) and amino acids (L-Serine, L-Proline, L-Valine, L-Threonine, L-Leucine, L-Isoleucine, L-Aspartic Acid, L-Lysine, L-Glutamic Acid, L-Methionine, L-Histidine, L-Phenylalanine, L-Arginine, L-Tyrosine, and L-Cystine) were quantified using external calibration to multi-point standard curves with R^2^ values > 0.99. MRM Targets are described in Supplementary Table S4, adapted from previous reports ^18,19^.

### Brewing Measurements

Specific gravity was measured using a PAL-1 Digital Refractometer (Atago), measured in Brix and converted to specific gravity.

